# Enhancement of Transgene Expression by NF-Y and CTCF

**DOI:** 10.1101/336156

**Authors:** Devon Zimmerman, Krupa Patel, Matthew Hall, Jacob Elmer

## Abstract

If a transgene is effectively delivered to a cell, its expression may still be limited by epigenetic mechanisms that silence the transgene. Indeed, once the transgene reaches the nucleus, it may be bound by histone proteins and condensed into heterochromatin or associated with repressor proteins that block transcription. In this study, we sought to enhance transgene expression by adding binding motifs for several different epigenetic enzymes either upstream or downstream of two promoters (CMV and EF1α). Screening these plasmids revealed that luciferase expression was enhanced 10-fold by the addition of a CCAAT box just upstream of the EF1α promoter to recruit nuclear transcription factor Y (NF-Y), while inserting a CCCTC-binding factor (CTCF) motif downstream of the EF1α promoter enhanced expression 14-fold (14.03 ± 6.54). ChIP assays confirmed that NF-Y and CTCF bound to the motifs that were added to each plasmid, but the presence of NF-Y and CTCF did not significantly affect the levels of histone acetylation (H3K9ac). Overall, these result show that transgene expression from the EF1α promoter can be significantly increased with motifs that recruit NF-Y or CTCF.

## Introduction

Condensation of DNA by histone proteins within the nucleus allows the cell to maintain a large amount of DNA (∼6 billion bp) within a relatively small nucleus (1-10 µm), but it can also sequester genes into inaccessible regions of chromatin (i.e. heterochromatin) that are transcriptionally inactive. However, specific post-translational modifications of histone residues can activate genes by either relaxing the local chromatin structure or recruiting transcription factors that enhance gene expression.^1,2^ For example, acetylation of the amine on lysine 9 of histone 3 (H3K9ac) activates genes by weakening the electrostatic interactions between the histone and DNA. In addition to acetylation, there are several other histone modifications and specific combinations thereof that can either activate or silence genes. Indeed, while H3K9ac is associated with transcriptional activation, trimethylation of that same residue (H3K9me3) is associated with transcriptional inactivation.^3^ Altogether, these mechanisms that control the transitions between tightly packed heterochromatin and relaxed euchromatin to control gene expression are known as epigenetics.

In addition to regulating host cell gene expression, epigenetic modifications can also defend the cell by silencing foreign viral genes.^4^ Unfortunately, both viral and non-viral gene therapy treatments can be hindered by these epigenetic defenses as well. For example, transgenes delivered via adenovirus, lentivirus, and γ-retrovirus have been silenced through methylation of the viral promoter^5,6^ and histone tail modifications (e.g. deacetylation of H3 and H4^7^ and/or the conversion of H3K9ac to H3K9me2^8^). Plasmid DNA delivered with non-viral vehicles can also be silenced by several different histone modifications, including H3K9me2, H3K9me3, H4K20me2, H4K20me3, and H3K27me3.^9–11^ Altogether, these previous studies show that epigenetic silencing mechanisms must be considered when performing gene delivery experiments.

A few strategies have been developed to reduce transgene silencing, including the inhibition of key epigenetic enzymes. For example, inhibition of lysine specific demethylase 1 (LSD1) with SL11144 can increase methylation of H3K4, decrease methylation of H3K9, and demethylate DNA methyltransferase 1 (DNMT1) to globally reduce DNA methylation and prevent gene silencing.^12,13^ Many DNMT inhibitors such as 5-Aza-2’-deoxycytidine and zebularine have been used to prevent methylation of DNA.^14,15^ Several inhibitors of histone deacetylases (HDACs) have also been shown to increase transgene expression, including Entinostat (a HDAC 1/3 inhibitor) ^16^, Tubacin (HDAC6i)^17^, Trichostatin A (HDAC6i)^18–20^, and Vorinostat (pan-HDACi)^21–23^. However, it is important to note that some of these HDAC inhibitors enhance transgene expression by influencing cytoplasmic transport instead of directly influencing histone modifications.

As an alternative to small molecule inhibitors, Kay et al. modified the sequence of the plasmid itself to avoid epigenetic silencing in a more localized fashion. Specifically, since the antibiotic resistance gene and origin of replication usually lie dormant within eukaryotic cells and serve as a nucleation point for heterochromatin formation that eventually spreads to the transgene, those elements were moved into an intron that is continuously transcribed. The resulting mini-intronic plasmid (MIP) significantly increases both the magnitude and duration of transgene expression i*n vitro* and *in vivo.*^11,24,25^ In addition, since promoter methylation can lead to gene silencing,^26,27^ removal of CpG motifs within CMV promoters has also been shown to provide higher long-term transgene expression.^28^

The goal of this work was to prevent transgene silencing by adding motifs for a variety of proteins that are known to epigenetically regulate gene expression. For example, a CCAAT motif was inserted upstream and downstream of two promoters (EF1α and CMV) to recruit nuclear factor Y (NF-Y), a histone-fold domain protein that is associated with transcriptional activation.^29^ Additional transcription factor binding sites (TFBS) that were inserted into the plasmid recruit other enzymes that provide a variety of modifications, including histone acetylation (ATF2^30^, SP1^31^, HNF4^32^, Zta^29^, POU6F1^33^, VSX2^33^), histone methylation (AP-1^34^ and AP-2a^34^, MYB^34^, NF-Y^34^, GFY-Staf^34^, Ying-Yang^35^), and general chromatin remodeling (Androgen Receptor^36^, CTCF^35^, Poly (dA:dT)^37^, FOXA1^38^). Overall, we show that adding NF-Y and CTCF motifs can significantly improve transgene expression, but in a location and promoter-specific fashion.

## Materials and Methods

### Preparation of Plasmids

The pEF-Luc plasmid was constructed by cloning a luciferase gene into the pEF-GFP plasmid (Plasmid #11154, Addgene, Cambridge, MA). The pCMV-Luc plasmid was then created by replacing the EF1α promoter with the CMV promoter. The sequence of each promoter and the corresponding plasmid backbone can be found in Figures S1, S2, and S3.

Each of the motifs shown in Table S1 were inserted 19 bp upstream of each promoter (at position −753 relative to the TSS of CMV and −222 for EF1α) using site directed mutagenesis, while the downstream motifs were inserted via oligo annealing cloning between EcoRI and KpnI sites (at position +13 relative to the TSS of CMV and +1194 for EF1α, see Figure 1). The sequence of each inserted motif, along with the corresponding promoter and reporter gene, was verified via Sanger sequencing.

**Figure 1:**
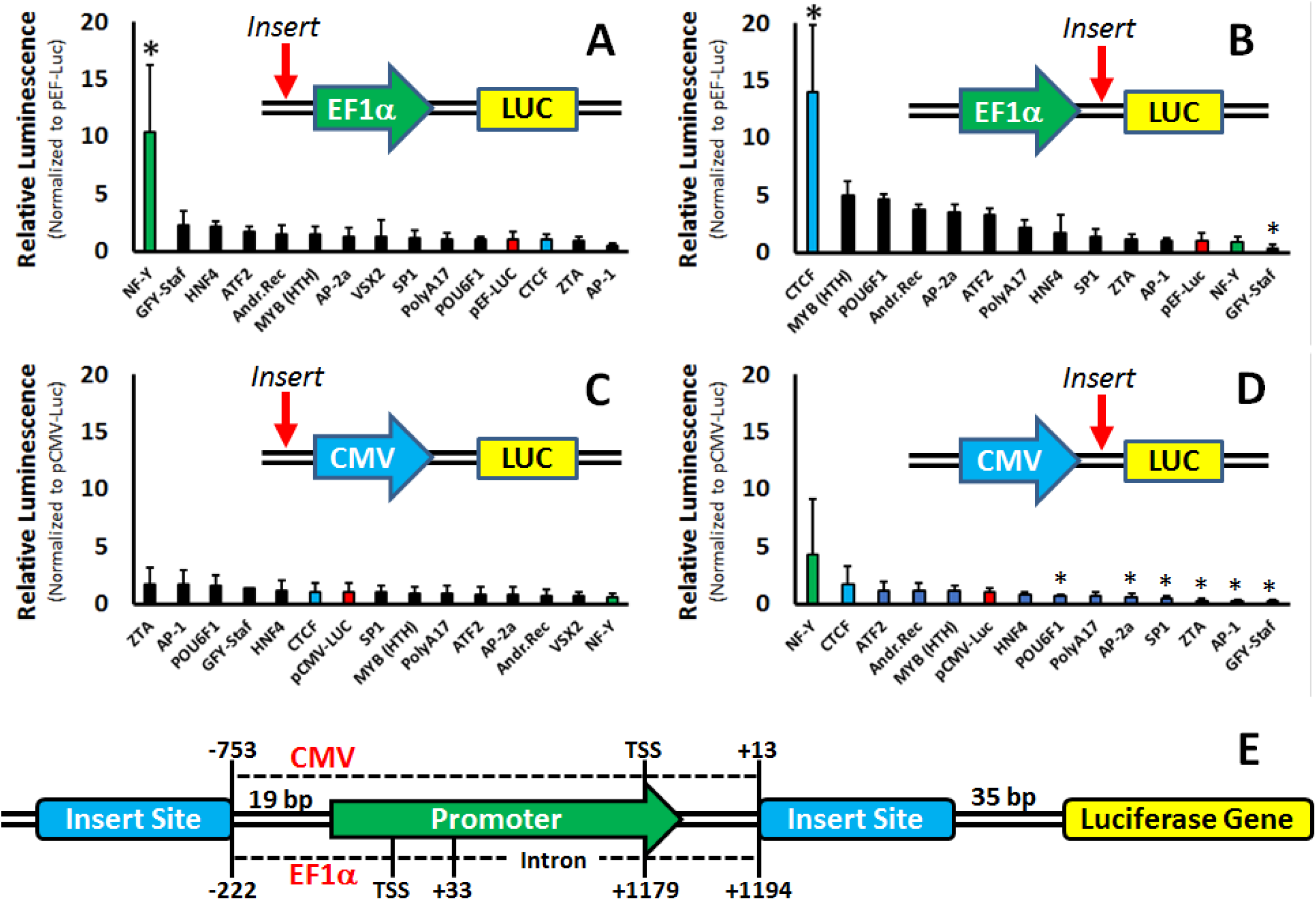
Effects of adding motifs upstream and downstream of the human EF1α (top panels, A & B) and viral CMV (bottom panels, C & D) promoters. The exact location of each insert relative to the transcription start site (TSS) is shown in the bottom panel (E). Experiments were performed in PC-3 (human prostate cancer) cells using branched PEI (8:1 N/P ratio).*significant difference relative to control plasmid (p ≤ 0.05 for n ≥ 3 independent experiments).

### Cell Transfections and Luciferase Assays

Human prostate cancer cells (PC-3) were seeded on 24 well plates at a density of 50,000 cells/well in fetal bovine serum-containing media (SCM) - Gibco^®^ RPMI 1640 (ThermoFisher Scientific Inc., Waltham, MA) - 24 hours prior to transfection. Polyplexes were prepared by mixing branched PEI (MW=25,000, Sigma Aldrich, St. Louis, MO) and the expression plasmid(s) in a 1:1 w/w ratio (8:1 N:P) with a total of 200 ng DNA/well, then incubating the mixture for 20 minutes at room temperature. Meanwhile, the SCM in each well was aspirated and replaced with serum-free media (SFM). Polyplexes were simultaneously added to each well and cells were then incubated for an additional 6 hours at 37°C in SFM, after which time the media was exchanged again with fresh SCM. Alternatively, to demonstrate the ability of the motifs to enhance transgene expression independently of the vehicle used (Figure 2) and ensure higher signals for ChIP assays (Figure 5), Lipofectamine LTX with PLUS reagent (Thermo Fisher, Hercules, CA) was used to transfect cells according to the manufacturer’s protocol. Finally, the data in Figures S5 and 1 were obtained by transfecting cells with jetPEI (Polyplus-transfection^®^, Illkirch, France) in a 5:1 nitrogen:phosphate (N:P) ratio.

**Figure 2:**
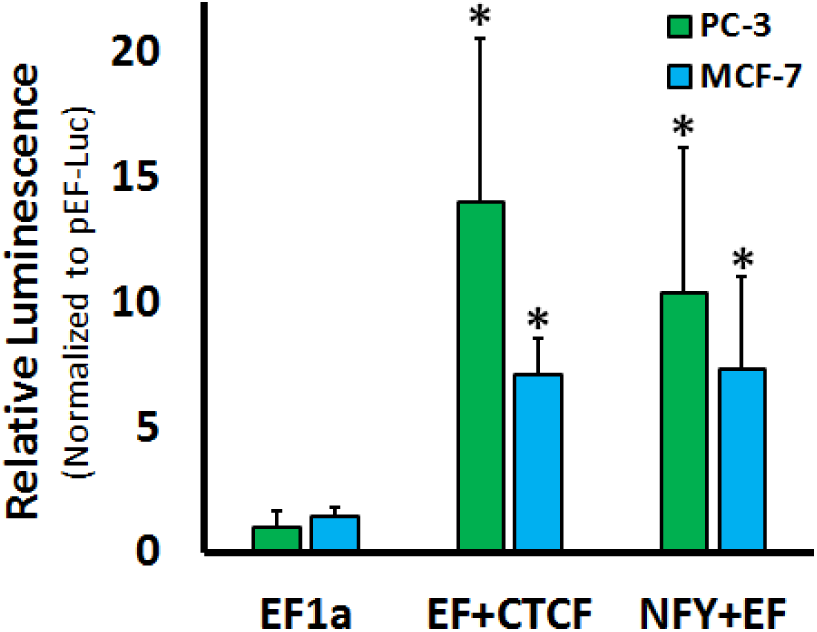
Effects of CTCF and NF-Y motifs on transgene expression in PC-3 and MCF-7 cells. PC-3 cells were transfected with PEI, while MCF-7 cells were transfected with Lipofectamine. Asterisks (*) indicate significant increases relative to EF1α (p ≤ 0.05 for n ≥ 3 independent experiments).

At 48 hours post-transfection, luciferase expression was measured using a Luciferase Assay kit (Promega, Madison, WI). Raw luminescence values for each triplicate were averaged and then normalized to the luminescence of the corresponding pEF-Luc or pCMV-Luc control without inserted motifs to obtain the relative luminescence values shown in each figure.

### Transient ChIP and qPCR Analysis

PC-3 cells were grown in T-75 flasks to ∼50-70% confluency, transfected with 1.5 µg of plasmid DNA using Lipofectamine^®^ LTX with PLUS^TM^ Reagent (ThermoFisher Scientific Inc., Waltham, MA), and then incubated for 48 hours at 37°C. Cells were then trypsinized to detach them from the flask and counted on a hemocytometer to determine their concentration. A sample of approximately 1-5×10^6^ cells was then treated with formaldehyde (1% final concentration) at room temperature for 10 minutes to cross-link the genomic DNA (including plasmids) to any bound proteins. Input samples of the genomic DNA were set aside for qPCR analysis, then chromatin immunoprecipitation was performed according to the manufacturer’s instructions for the SimpleChIP^®^ Enzymatic Chromatin IP Kit (#9003, Cell Signaling Technology, Danvers, MA) with minor alterations. Specifically, cells were lysed via sonication with four 20-second pulses at 10% power with a 1 minute incubation between pulses with a Branson Digital Sonifier and a Branson Model 102C Probe. With the exception of the NF-YA antibody (Abcam, #ab139402), all antibodies used for immunoprecipitations were obtained from Cell Signaling Technologies (#3418S anti-CTCF Ab, #9649S anti-H3K9ac Ab; IgG & anti-H3 antibodies were included in the SimpleChIP^®^ kit). Threshold cycle (C_T_) values were then measured for each sample using plasmid-specific primers and SYBR^TM^ Select Master Mix for CFX (ThermoFisher Scientific Inc., Waltham, MA) on a QuantStudio3 Real Time PCR System. The percent input (%input) for each sample was then calculated with Equation 1.

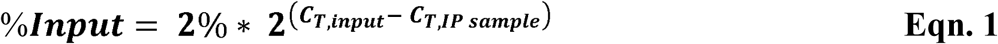

### Plasmid Copy Number

The fold-change in nuclear uptake of lead enhancer plasmids relative to the control plasmid was quantified by determining the copy number of transfected pDNA relative to the actively transcribed housekeeping gene RPL30. qPCR with SYBR^®^ Green was performed using 2 μl SimpleChIP^®^ Human RPL30 Exon 3 Primers from Cell Signaling Technology and 1 μl each of forward and reverse primers specific to the pDNA enhancer sequence. Relative copy number was determined using the 2^ΔCt^ method where ΔC_t_ is the difference in C_t_ between reactions performed with RPL30 primers and enhancer primers. The fold-change in copy number was determined using the 2^ΔΔCt^ method in which ΔΔC_t_ is the difference in ΔC_t_ between the sample plasmids and the control plasmid.

### Statistics

Statistical significance was evaluated in R studio using Analysis of Variation testing (ANOVA). For all experiments, a 95% confidence interval test was used to determine if data was statistically significant (p<0.05).

## Results and Discussion

### Effects of Motifs on Transgene Expression

Of the 14 motifs tested, only a few significantly enhanced transgene expression (Figure 1; see Table S1 for numerical values). Some of the strongest enhancement was observed with the NF-Y motif (TCAG<u>CCAATC</u>AGCGAG), which significantly increased transgene (luciferase) expression when inserted upstream of the EF1α promoter (10.4 ± 5.8-fold). Inserting the NF-Y motif downstream of the CMV promoter also seemed to increase transgene expression about 4-fold, but the effect was not statistically significant. Likewise, no enhancement was observed when the NF-Y motif was inserted upstream of the CMV promoter. This lack of enhancement of the CMV promoter may be due to the fact that eight separate NF-Y motifs (CCAAT or ATTGG, see Figure S2) are distributed throughout the native CMV promoter, such that an additional CCAAT site may be redundant. In contrast, the EF1α promoter does not contain any CCAAT motifs.

Strong enhancement (14.0 ± 6.5-fold) was also observed when the motif for CCCTC-binding factor (CTCF, AGACCACCAGAGGGCACCA) was inserted downstream of the EF1α promoter. As was the case for the NF-Y motif, the CTCF motif did not enhance expression from the CMV promoter. Interestingly, further analysis revealed that the CMV promoter already contains a high-affinity CTCF motif (Figure S4), while the EF1α promoter contains only a partial low-affinity CTCF motif. It is also worth mentioning that the plasmid backbone itself contains three high-affinity CTCF sites in the PolyA region and another high-affinity CTCF site just upstream of the EF1α/CMV promoters (Figure S4). Therefore, an additional CTCF site may be redundant in the CMV promoter and upstream of the EF1α promoter, but beneficial when inserted downstream of the EF1α promoter.

It is also interesting to note that a directly comparison of the two native promoters shows that the CMV promoter (which contains NF-Y and CTCF motifs) consistently provides 4-fold higher transgene expression than the EF1α promoter (Figure S7). However, as shown in Figure 1, adding either motif to the EF1α promoter provides 2-3 times more transgene expression than the CMV promoter. We also attempted to further enhance expression by creating an EF1α promoter with both an upstream NF-Y motif and a downstream CTCF motif, but the combination of motifs did not significantly increase luciferase expression relative to an EF1α promoter with a single NF-Y motif. (Figure S5).

In contrast to NF-Y and CTCF, some of the motifs significantly decreased transgene expression when inserted downstream of the promoters. For example, inserting the GFY-Staf motif downstream of both promoters decreased luciferase expression by 60-80%, while several other motifs (POU6F1, AP-1, AP-2a, Sp1, and Zta) significantly decreased expression (30-80%) from the CMV promoter.

To determine if the enhancement observed with the NF-Y and CTCF motifs could also be observed with other cell lines and other gene delivery vehicles, we used Lipofectamine to transfect the plasmids into breast cancer cells (MCF-7). As shown in Figure 2, plasmids containing the upstream NF-Y or CTCF motifs significantly enhanced luciferase expression in the MCF-7 cell line, although the magnitude of enhancement was only 7-fold. Similar experiments were also conducted with the Jurkat line of leukemia T cells, but the motifs did not provide any significant enhancement relative to the native EF1α promoter (data not shown).

### Further Investigation of the NF-Y Motif

Nuclear Transcription Factor Y (NF-Y), also known as CCAAT Binding Factor (CBF) or CCAAT Protein 1 (CP1), is a trimeric protein comprised of three subunits: NF-YA, NF-YB, and NF-YC.^39^ While the NF-YA subunit is responsible for binding the CCAAT motif, the NF-YB and NF-YC subunits contain histone-fold domains (HFDs) that can mimic and replace histones H2A & H2B, respectively.^40^ NF-Y binding can even occur in relatively inaccessible regions of heterochromatin, thereby allowing other transcription factors and RNA Polymerase to activate transcription of otherwise silenced genes.^41^ NF-Y is also associated with several histone modifications (H3K4me3, H3K79me2, and H3K36me3) that activate gene expression.^42^ In addition, the NF-YB subunit can also be post-translationally modified at Lys138 to a ubiquitinated form that resembles H2B Lys120 monoubiquitination, which is associated with transcriptional activation.^40^ Therefore, there are several ways in which adding an NF-Y motif to the EF1α promoter could provide the observed enhancement of transgene expression.

In the human genome, NF-Y motifs are frequently found in pairs with short spacers of 24-31 bp between each CCAAT box.^39,42,43^ Since the CCAAT motif identified in our initial screen only contains a single CCAAT box, we investigated the effects of inserting tandem CCAAT motifs separated by 24, 29, or 31 bp spacers upstream of the EF1α promoter. Surprisingly, each of the tandem NF-Y motifs significantly decreased transgene expression by 50-60% relative to the single NF-Y motif (Figure S6).

While tandem NF-Y sites are common in the human genome, some of the individual NF-Y motifs in the CMV promoter are separated by approximately 200 bp, which roughly corresponds to the distance between nucleosomes. We next investigated if nucleosomal spacing of NF-Y motifs could further enhance transgene expression by inserting additional NF-Y motifs at other sites in the plasmid (Figure 3). First of all, it is interesting to note that moving the single NF-Y to the ampicillin resistance gene (position −471), inside the EF1α promoter (position −45), or even past the TSS to the intron within the EF1α promoter region (position +145) provided the same level of expression obtained with the CCAAT motif at its initial location (position −248). In most cases, combining pairs of NF-Y motifs at these locations had no significant effect on transgene expression. Introducing pairs of NF-Y motifs gave no significant increase in transgene expression, although a slight (µ4-fold) increase in transgene expression seemed to occur when a combination of NF-Y motifs was inserted at −471 and −248 (but the overall difference was not significant).

**Figure 3:**
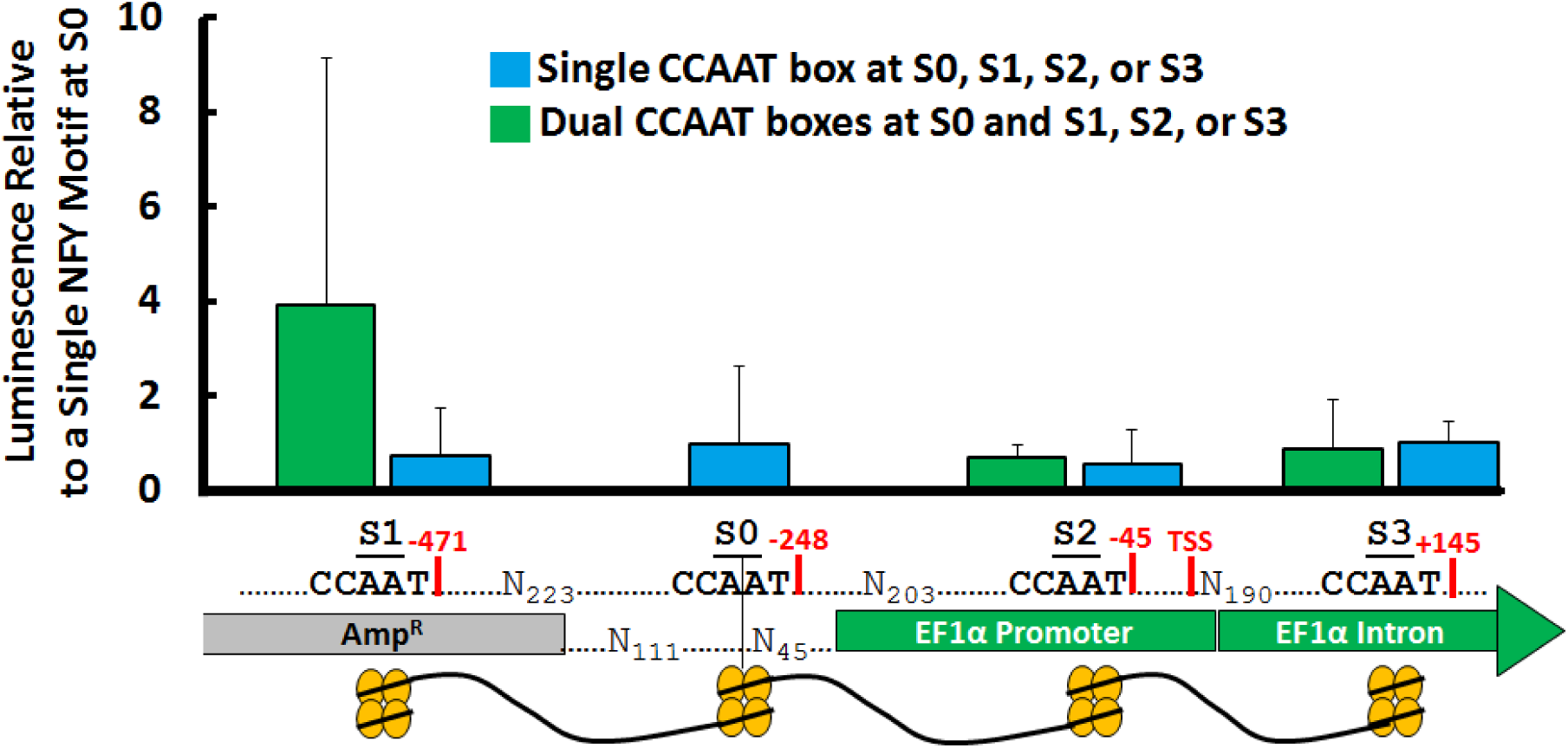
Effects of single NF-Y motifs and pairs of NF-Y motifs relative to a plasmid with a single NF-Y motif at position −248 relative to the transcription start site (TSS) of the EF1α promoter. The S0 site is located 19 bp upstream of the EF1α promoter region and was used for the initial motif screening experiments. The blue bars represent plasmids with single NF-Y motifs inserted at each location, while the green bars represent plasmids with NF-Y motifs present at that position and position −248.

### Further Investigation of the CTCF Motif

CTCF has a wide range of functions within the genome, including both repression and activation of genes.^44^ Regarding gene activation, the nucleosomes that flank CTCF sites tend to be highly enriched in the histone variant H2A.Z, which flanks nucleosome free regions and is associated with activating histone modifications (e.g., H3K4me3 and H3K4me).^45–47^ The histone variant H2A.Z itself is multifunctional protein involved both with transcriptional inactivation and activation through recruitment of pioneering transcription factors such as FOXA2.^48^ CTCF has also been shown to activate poly(ADP-ribose) polymerase 1 (PARP1) which ADP-ribosylates DNMT1, thus preventing the enzyme from methylating CpG motifs.^49,50^ Recently, CTCF has also been shown to control chromatin conformation by interacting with cohesin. Cohesin, a trimeric protein comprised of SMC1, SMC3, and SCC3 subunits,^51^ does not directly bind to the DNA but instead interacts with CTCF through its SCC3 subunit.^52^ The CTCF/cohesion complex forms loops of chromatin between oppositely oriented CTCF motifs, thereby bringing distal enhancer elements in contact with promoters to increase transcription.^53,54^

Previous studies have shown that CTCF binds DNA using 11 zinc finger domains.^55^ The number of zinc fingers that bind the target DNA and the orientation of the binding site (plus/minus) have also been shown to determine the effects of CTCF binding (e.g., activation or repression of a gene). Since the CTCF motif used in our initial screen only bound 4 of the 11 zinc fingers in the sense (plus) orientation, we attempted to stabilize CTCF binding and potentially increase transgene expression by testing two additional CTCF motifs: (1) an alternative CTCF motif with a slightly different sequence than the initial CTCF motif35 and (2) a longer CTCF motif that binds 8 of the 11 CTCF zinc finger domains.^55^ Both of these sequences were tested in the plus and minus orientations as well. Figure 5 shows that the alternative CTCF motif provided significantly decreased transgene expression, but elongating the CTCF motif to bind additional zinc finger domains within CTCF did significantly increase transgene expression 3-fold relative to the initial CTCF motif. Interestingly, however, switching the orientation of the longer CTCF motif from minus to plus caused a significant decrease in transgene expression relative to the initial CTCF motif.

### Epigenetic Effects of the NF-Y and CTCF Motifs

ChIP assays were performed to confirm that the enhancement observed with the NF-Y and CTCF motifs was indeed due to NF-Y or CTCF binding the plasmid. As shown in Figure 4a, the amount of NF-Y bound to the EF1α promoter region significantly increased upon addition of the NF-Y motif to the EF1α promoter. Likewise, inserting the expanded CTCF motif with a stabilizer domain (see Figure 4) downstream of the EF1α promoter also exhibited a significant increase in CTCF binding relative to the native EF1α promoter. In both cases, the %input values for the NF-Y and CTCF samples were also significantly higher than the %input values for their corresponding negative control IgG. Therefore, it appears that the enhancement observed with these motifs is due to the recruitment of NF-Y or CTCF to the plasmid. However, the ChIP experiments did not show any changes in histone H3 binding or local H3K9 acetylation, so the change in transgene expression may be due to some other type of histone modification or the recruitment of additional transcription factors/proteins by NF-Y or CTCF.

**Figure 4:**
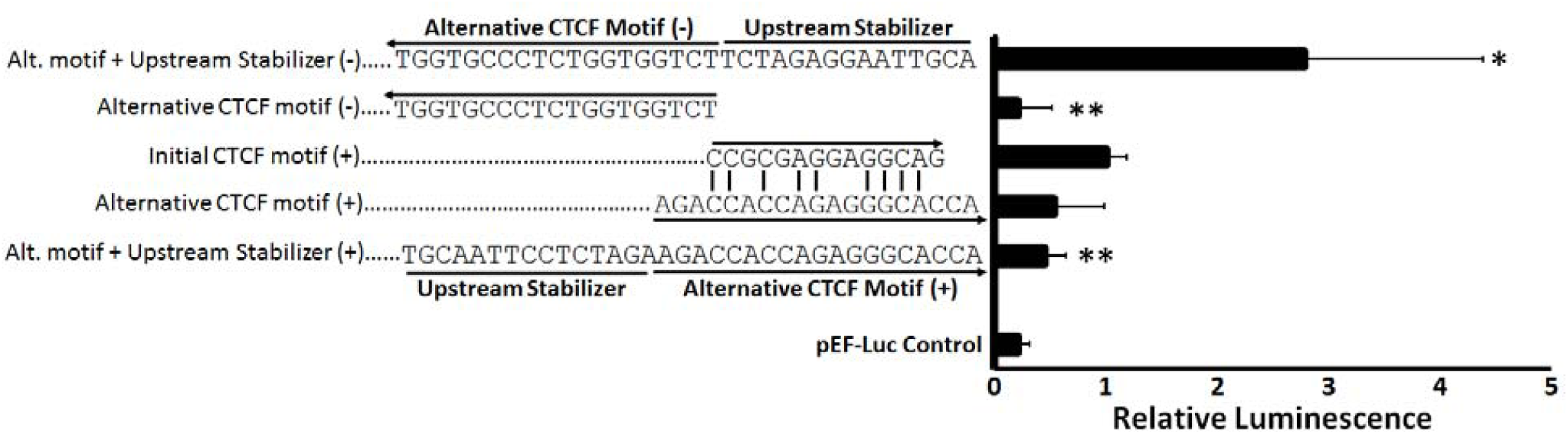
Optimization of the CTCF sequence and its orientation. The initial CTCF motif was used in the screening experiments, while the alternative CTCF motif contains a slightly different sequence that binds 4 the 11 zinc finger domains within CTCF. In addition, an upstream stabilizer sequence was also added to the core CTCF motif to facilitate binding of 4 more zinc finger domains from CTCF. Each of these motifs was inserted downstream of the EF1α promoter in both the sense (+) and antisense (-) orientation. Single asterisks (*) indicate plasmids with transgene expression/luminescence values significantly higher than plasmids containing the initial CTCF, while double asterisks (**) indicate significant decreases in luminescence (p <0.05, n ≥ 3 independent experiments).

**Figure 5:**
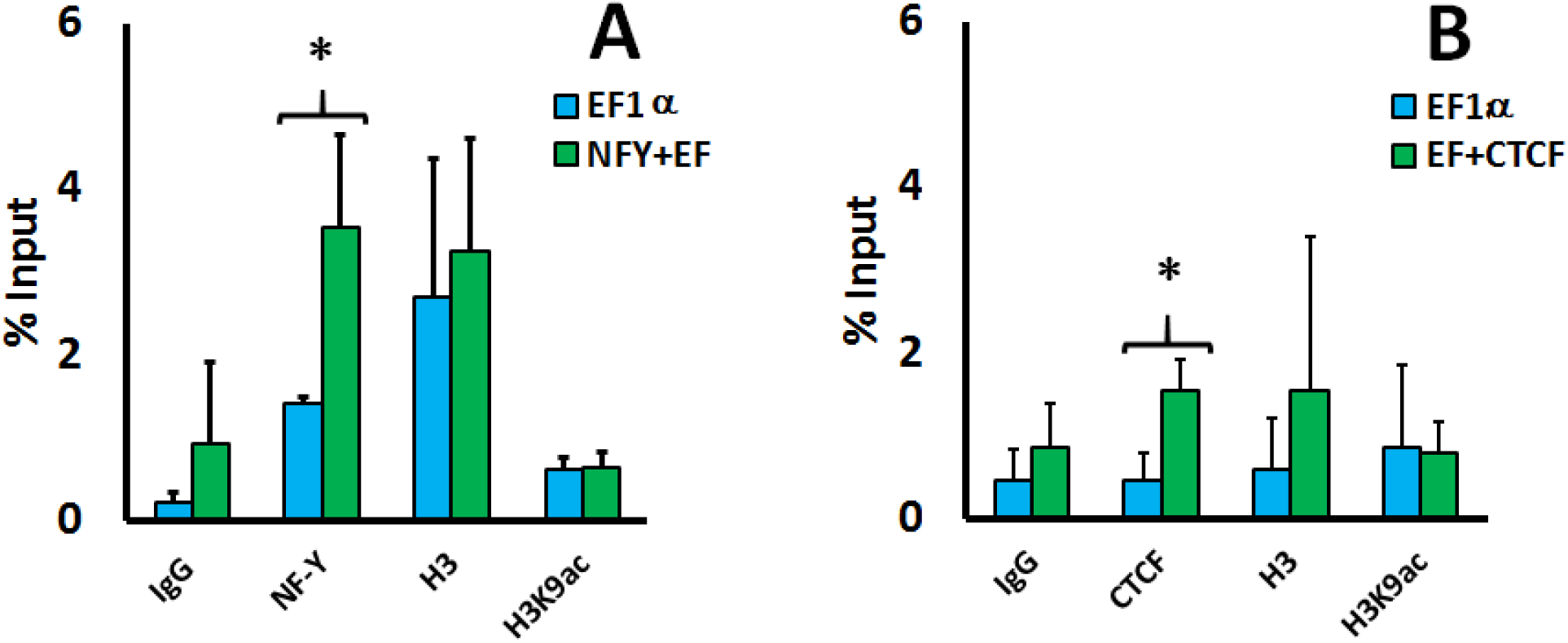
–Transiently transfected plasmids with the NF-Y motif (left panel, A) or the CTCF motif (right panel, B) were tested for NF-Y, CTCF, H3, and H3K9ac binding relative to the native EF1α promoter in PC-3 cells. Asterisks (*) indicate a significant increase in occupancy of the target protein relative to the native promoter (EF1α).

### Nuclear Uptake Effects

Since it is possible that the enhancement observed with each motif may be due to NF-Y or CTCF binding the plasmid in the cytoplasm and then increasing its rate of nuclear transport, we also calculated the fold-change in nuclear copy number for the plasmids with the motifs (plasmid copy numbers were normalized to the housekeeping gene RPL30). Interestingly, Figure 6 shows that the addition of the NF-Y motif significantly decreases the plasmid copy number by approximately 20% (relative to a plasmid containing the native EF1α promoter). In contrast, the CTCF motif had no significant effect on plasmid copy number. Therefore, it appears that the enhancement provided by the NF-Y and CTCF motifs are not due to an increase in nuclear copy number.

**Figure 6:**
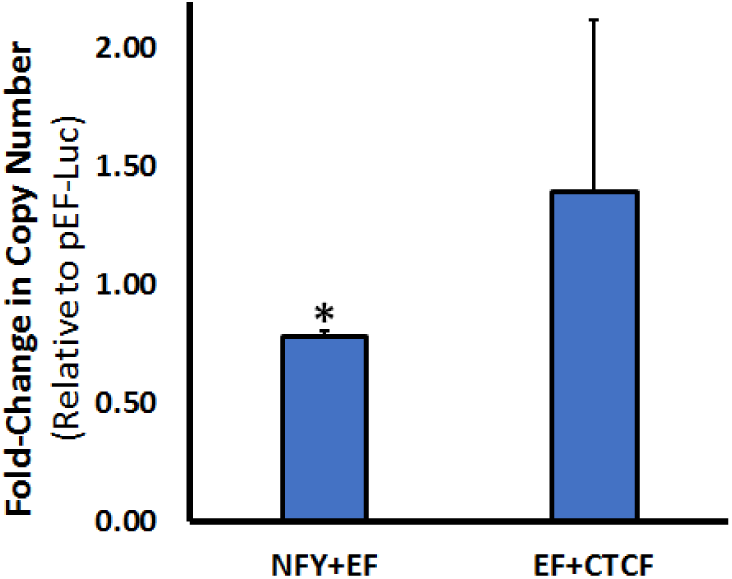
Fold-change in copy number of plasmids with the NF-Y motif (NFY+EF) or the CTCF motif (EF+CTCF) relative to a plasmid with the native EF1α promoter. Copy numbers were normalized to the housekeeping gene RPL30 (data not shown). Asterisks (*) indicate a significant decrease in nuclear copy number relative to the plasmid with the native promoter (EF1α).

## Conclusion

Previous studies have shown that foreign transgenes are susceptible to epigenetic silencing, but our results show that transgene expression can be significantly increased by modifying the EF1α promoter to recruit pioneering transcription factors like NF-Y and chromatin remodeling enzymes like CTCF. The exact mechanism by which NF-Y and CTCF increase transgene expression is currently unknown, but both of these proteins are known to significantly remodel chromatin by replacing histones (NF-Y) or forming chromatin loops that insulate specific loci (CTCF). Therefore, the conformation of the plasmid chromatin may be a particularly important epigenetic factor that regulates transgene expression.

## Acknowledgements

This work was supported by NSF CBET Grant 1403214. The authors have no conflicts of interest related to this work.

